# Cryptic viruses shape trophic networks after biological invasions

**DOI:** 10.64898/2026.11.16.766434

**Authors:** J. Pérez-Rodríguez, L. Hernández-Pelegrín, R. García-García, E. Martín-Sarrate, A. Tena, L. Gasmi, S. Herrero

**Author notes:** Corresponding authors: Jesica Pérez-Rodríguez, Laila Gasmi and Salvador Herrero, University Institute of Biotechnology and Biomedicine (BIOTECMED), Universitat de València, Unidad Mixta Gestión Biotecnológica de Plagas UV-IVIA, Dr Moliner 50, 46100 Burjassot, Valencia, Spain.

## Abstract

Biological invasions restructure ecological communities by bringing previously isolated organisms into contact, but the viral flow within the trophic network and its consequences have not been established. Here, we show viral exchange following the introduction of a parasitoid wasp released for classical biological control in citrus. Within a few years, viruses spread bidirectionally between native and exotic host-parasitoid trophic chains and reached a generalist hyperparasitoid at the fourth trophic level. Cohabitation and multiparasitism, two ubiquitous interactions between the two trophic chains, were sufficient for viruses to be transferred between species and maintained transgenerationally. A virus acquired by the introduced parasitoid increased its offspring production by 30%. Our results identify cryptic viruses as an overlooked dimension of biological invasions, reshaping trophic networks and the outcome of the species we deliberately introduce.

## Introduction

Arthropod species are relocated around the world at an unprecedented rate (*1–3*). Whether introduced accidentally or deliberately, arthropods carry an extensive community of associated microorganisms, including viruses (*4*, *5*). Sequencing surveys have reported large numbers of RNA viruses that cause no obvious symptoms in their arthropod host (*6*, *7*). These cryptic viruses establish covert infections and are transmitted both vertically and horizontally, yet in the few cases examined in detail they alter the physiology, behavior, and reproductive success of their hosts (*8–10*).

Once established, both accidentally and deliberately introduced arthropods become embedded in the native trophic network and establish interactions with native species of their and other trophic chains (*11*, *12*). Each of these interactions is a potential route for viral exchange in both directions, as viruses carried by the newcomer may enter the resident community and resident viruses may be acquired by the newcomer (*13*). Such exchange has long been expected to reshape species interactions and community composition (*14*). Virus sharing between introduced and native species has been reported in plants (*15*, *16*) and animals (*17*, *18*), but transmission has been inferred rather than demonstrated. Direct evidence that viruses flow between exotic and native species across trophic levels is lacking, and the ecological consequences of such spillover remain largely unknown.

Classical biological control programs provide a perfect scenario to study the viral flow among exotic and native species. These programs involve the deliberate introduction of living organisms to suppress an invasive pest (*19*, *20*). Classical biological control is prioritized in current pest management programs because it offers an environmentally safe, targeted, and sustainable alternative to chemical pesticides (*21*, *22*). Herein, we used a program carried out in Spanish citrus that have long been affected by a mealybug pest, *Planococcus citri* (Risso) (Hemiptera: Pseudococcidae), that can be naturally controlled by the native parasitoid wasp, *Anagyrus vladimiri* Triapitsyn (Hymenoptera: Encyrtidae) (*23*, *24*); hereinafter native chain. In 2009, an invasive mealybug, *Delottococcus aberiae* (De Lotto) (Hemiptera: Pseudococcidae), arrived from South Africa and spread rapidly, causing significant economic losses (*25*). Its South African parasitoid wasp *Anagyrus aberiae* Guerrieri (Hymenoptera: Encyrtidae) was imported and released from 2020 onwards (*26*); hereinafter exotic chain. The two trophic chains now overlap on the same trees, where the parasitoid wasps of both chains are attacked by the same generalist hyperparasitoid, *Chartocerus* sp. (Hymenoptera: Signiphoridae), which develops on them and thus occupies a fourth trophic level (Fig 1A). This scenario provided an ideal study system where two phylogenetically related parasitoids and their respective hosts, that were long geographically separated, co-occur, allowing new host-virus interactions that might modulate the ecosystem.

**Figure 1.**
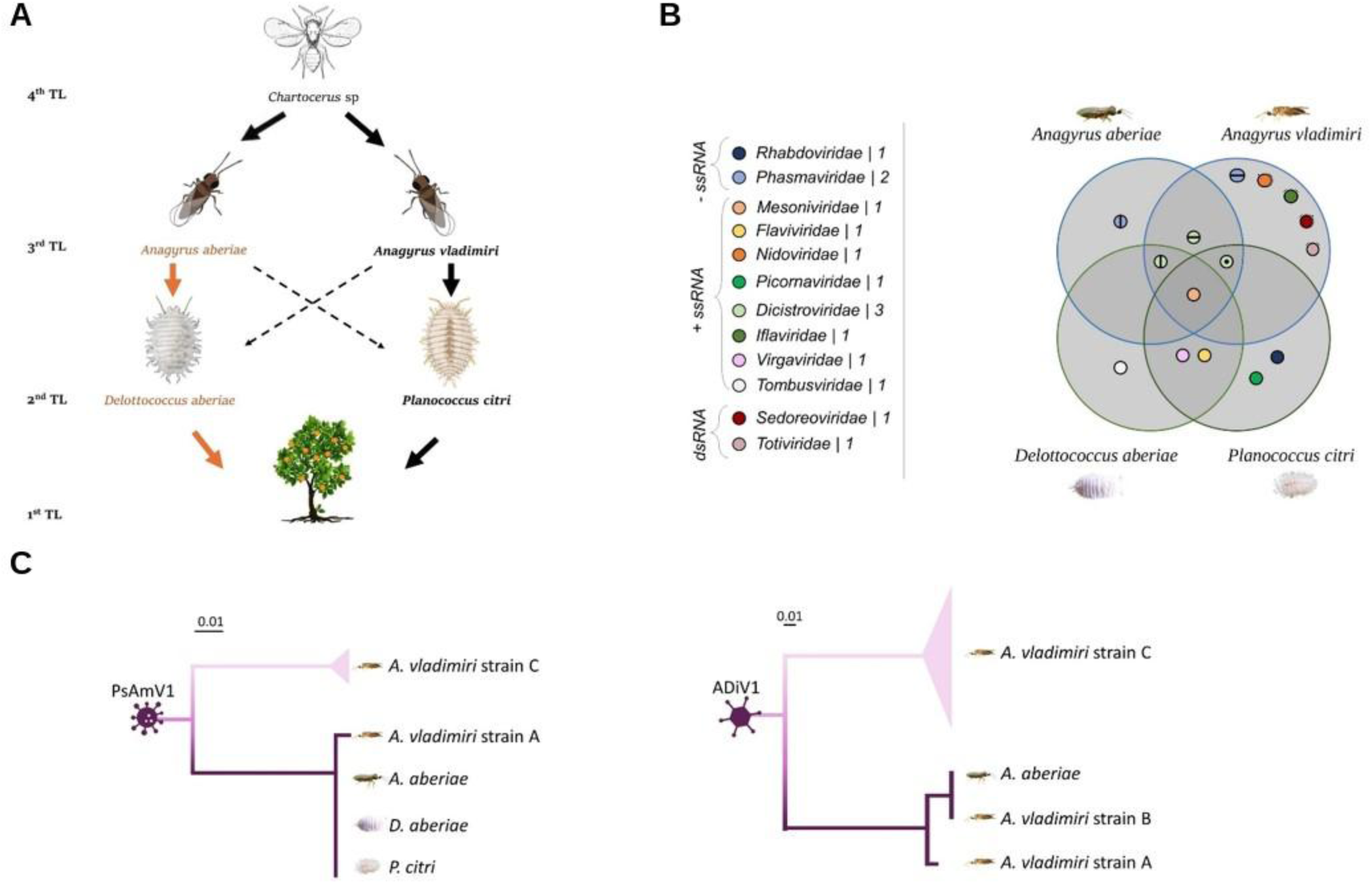
Rich viral biodiversity in the native and exotic *Anagyrus*-mealybug trophic chains. **(A)** Schematic overview of the studied trophic network, illustrating different trophic levels: first trophic level (1^st^ TL): citrus plant, second trophic level (2^nd^ TL): pseudococcid mealybugs, third trophic level (3^rd^): primary parasitoids, and fourth trophic level (4^th^ TL): hyperparasitoids. The naturalized system is depicted in black letters and in orange color the invasive pest and its exotic introduced natural enemy. Dashed lines represent the possibility of cross-oviposition in non-permissive host. **(B)** Venn diagram representing the viral families found in *P. citri*, *D. aberiae*, *A. vladimiri*, and *A. aberiae*. **(C)** Phylogenetic inference of viral genomes obtained from diverse samples of the Pseudococcidae*–Anagyrus* trophic network. Phylogenetic distances are indicated above each tree.

Here, we first demonstrate that viruses flow, in both directions, between exotic and native chains within a few years of contact in the field, crossing trophic levels and reaching hyperparasitoids that attack both. Following these findings, we identify two possible mechanisms for this spontaneous transmission between chains and show that newly acquired viruses are inherited across generations, conferring measurable advantages to the new host.

## Material and Methods

### Insect colonies

Colonies of *Delottococcus aberiae* were obtained from the Instituto Valenciano de Investigaciones Agrarias (IVIA; Moncada, Spain) facilities and continuously reared on organic lemons (*Citrus limon* (L.) Osbeck), at the University of Valencia. Colonies were kept inside cardboard boxes (30 × 22 × 25 cm), placed in a dark climatic chamber at 25 ± 1 °C and 70 ± 5% relative humidity (RH).

*Planococcus citri* was obtained as first instar nymphs from the State Insectary of Silla (Valencia, Spain), where it was mass reared on butternut squash (*Cucurbita moschata* L. Duchesne) following the procedures described in (*27*). It was subsequently reared for more than five generations on green bean pods (*Phaseolus vulgaris* L.) at the University of Valencia facilities. Colonies were kept in plastic containers (30.5 × 24.5 × 20 cm) with a perforated lid (15 × 15 cm) on top, covered with fine mesh and maintained at 25 °C ± 1, 70 ± 5% RH and a 16:8 h light:dark (L:D) photoperiod.

*Anagyrus aberiae* parasitoids belonged to five different sources: (i) Isofemale lines. Seven isofemale lines were randomly selected from the 21 maintained at IVIA. Each line was established before the implementation of the classical biological control program against *D. aberiae*, originating from single unmated females at the time the parasitoids were imported from South Africa in 2020. Because unmated females produce only male offspring (arrhenotoky), each founding female was subsequently backcrossed with her own male offspring to obtain female progeny and establish the line. Since their establishment, these lines have been maintained in isolation, without exposure to any other insect species apart from their host *D. aberiae*. (ii) Line 17/19UV. A mixed line obtained by combining two randomly selected isofemale lines (17 and 19) in 2024, maintained thereafter as a single colony at the University of Valencia. (iii) Mass rearing. Individuals from mass rearing located at TRAGSA facilities (Moncada, Spain). This population was originally established by crossing the South African isofemale lines in 2020 and has been refreshed annually with field-collected individuals to preserve genetic diversity. (iv) Field-collected individuals. These parasitoids corresponded to the offspring of mass-released parasitoids in 2020. To obtain them, citrus twigs infested with *D. aberiae* were kept in plastic containers for three weeks and checked daily for parasitoids emergence. Once parasitoids emerged, they were isolated and identified under a binocular stereomicroscope. (v) *Anagyrus aberiae* laboratory population (SelUV) in which the *Anagyrus vladimiri* reovirus (AnvRV) had been introduced experimentally through cohabitation with *A. vladimiri*. The virus became established in this population but did not reach fixation, and its prevalence varied among individuals.

All *A. aberiae* individuals were maintained at laboratory conditions in separated Bugdorm® cages sized 24.5 x 24.5 x 24.5 cm, with a mesh size of 44 x 32 | 650 µm aperture and a sleeve opening of 16 cm diameter. Parasitoids were supplied with water through an embedded cotton ball and with 70:30 (v/v) honey–water droplets. Bugdorm® cages were kept in a climatic chamber at 25 ± 1°C, 70 ± 10% RH and 16:8 L:D regime.

*Anagyrus vladimiri* specimens were received as pupae from two commercial biological control companies (A and B) located in Spain. Upon arrival, they were introduced into separated Bugdorm® cages and maintained in the same conditions described for *A. aberiae*.

The hyperparasitoid *Chartocerus* sp., was field-collected with the same procedure as the one described for field-collected *A. aberiae*. As *Chartocerus* sp used in this study was a gregarious species, each specimen selected for processing was randomly chosen from different mealybug hosts, ensuring that they did not originate from the same egg clutch (pseudo replicates).

### High throughput sequencing and viral characterization

Viral discovery was performed using non-parasitized *D. aberiae* and *P. citri,* the parasitoids *A. aberiae* and *A. vladimiri,* and mealybug hosts parasitized by each parasitoid species under laboratory conditions. Parasitized mealybugs were obtained from laboratory colonies, whereas non-parasitized mealybugs comprised individuals from both laboratory colonies and field-collected in Benavites (Valencia, Spain; 39.7462° N, 0.2494° W, year 2023). Field-collected individuals were subsequently reared in the laboratory for more than two generations to ensure they were parasitoid-free. *Anagyrus aberiae* females were obtained from the mass-reared population and *A. vladimiri* females from the two biological control companies.

In the case of parasitized mealybugs, each *Anagyrus* species was allowed to oviposit in both permissive (the natural host) and non-permissive (the other pseudococcid species) mealybug species. For each parasitoid-mealybug combination, the viral diversity transmitted through oviposition was assessed and therefore five treatments were established: i) mealybugs parasitized for 6 h (hosting the parasitoid egg), ii) mealybugs parasitized for 24 h (hosting the parasitoid egg), iii) mealybugs parasitized for 36 h, after which the parasitoid egg was removed, iv) non-parasitized mealybugs, and v) naïve parasitoids. The parasitism assay was conducted in a glass Petri dish (5 cm diameter) with a piece of lemon infested with a mix of six third instar nymphs and six young females *D. aberiae* or 15 cm long green bean pods infested by the same number and instars of *P. citri.* Following, a 48h-72h old mated *Anagyrus* female was released in each assay plate and visual observations were conducted until parasitoids oviposited. Upon oviposition, each parasitoid was stored at –20°C and the parasitized mealybug was maintained 6, 24 or 36h depending on the treatment, on a piece of lemon or green bean pod, in a climatic chamber at 25 ± 1°C, 70±10% RH and 16: 8 L:D regime. At 6 and 24 hours post oviposition, the corresponding mealybugs were stored individually at –20°C. Mealybugs from the 36-hour treatment group were dissected using entomological pins to remove the parasitoid egg and then frozen at –20 °C. For each parasitoid-mealybug and time point combination, 16 independent replicates were conducted.

Total RNA from the different pooled samples were isolated using TRIZOL reagent (TRItidy G, ITW Reagents), following the manufacturer protocol. Library preparation and ribosomal RNA (rRNA) depletion were performed using the Truseq Total RNA library kit with Ribo-Zero (Illumina, San Diego, CA, USA). RNA sequencing was performed by using the NovaSeq 6000 platform rendering paired-end reads and a total of 40 million reads per sample. Library construction and sequencing services were performed at Macrogen (Seoul, Korea). Sequencing data are deposited in the Sequence Read Archive (SRA) of the National Centre of Biotechnology Information (NCBI) under the BioProject ID: PRJNA1193656 (Table S1).

The presence of RNA viruses was assessed in six *de novo* assembled transcriptomes derived from *A. aberiae, A. vladimiri*, *D. aberiae* (non-parasitized and parasitized), and *P. citri* (non-parasitized and parasitized) as previously described in (*28*). Briefly, the quality of the raw sequencing reads was analyzed using fastQC (https://github.com/s-andrews/FastQC). The adaptors and low-quality sequences were trimmed using Trimmomatic (*29*), and the clean reads were *de novo* assembled into contigs using Trinity v2.9.0 (*30*). Default parameters were used when no additional information is provided. To investigate their viral origin, the assembled contigs were mapped against a viral query using tblastn (protein query against translated nucleotide database) with a threshold value of 1 × 10^−5^. The viral query contained the GenBank protein sequences classified within the Riboviria domain (RNA viruses) and identified in invertebrate hosts. Putative viral contigs were further filtered to remove (a) contigs showing similarity to their respective insect reference genome after blastn (e-value > 10^-100^), (b) contigs smaller than 1000 bp in length, and (c) contigs containing no open reading frames (ORFs) after evaluation with ORFfinder (https://www.ncbi.nlm.nih.gov/orffinder/). The remaining contigs were defined as RNA viruses. The new viral genomes were characterized by annotating the length, position and function of each ORF using ORFfinder and the blastp search against the nr protein database on NCBI. Then, the new viruses were phylogenetically classified using the RNA-dependent RNA polymerase (RdRp) region. The RdRp protein sequences of the newly described viruses and their closest relatives (identified with blastp) were aligned using T-coffee mode “M-coffee” (*31*). The resulting alignments were manually trimmed with BioEdit to eliminate the poorly aligned sequences at the 5’ and 3’ends. The curated alignments were uploaded into IQtree2, which inferred the phylogenetic trees using the maximum-likelihood method, with default parameters, and 1000 ultrafast bootstraps (*32*). The resulting phylogenetic trees were visualized using iTOL (*33*).

### Characterizing viral flow across the trophic network

To investigate the transmission of viruses within the three trophic levels (mealybugs, parasitoids, and hyperparasitoids), we first analyzed the viral diversity and abundance in randomly selected *A. aberiae* individuals from different isolines. This enabled us to identify those viruses that are associated with the native range of *A. aberiae* and subsequently introduced to the native network following the importation of the parasitoid. A second subset of *A. aberiae* individuals were collected from Spanish citrus fields during surveys performed in 2024, four years following introduction and establishment of the parasitoid. In the same surveys, hyperparasitoid individuals of the genus *Chartocerus* were collected. Two years later (2026), and to assess the evolution of the viral flow in field populations, *A. aberiae* individuals were collected from the field. Sampling in the field was performed by cutting citrus twigs infested with *D. aberiae* which were kept in plastic containers for three weeks and checked daily for parasitoids emergence. Once emerged, primary parasitoids or hyperparasitoids were isolated and identified under a binocular stereomicroscope.

A total of 52 *A. aberiae* individuals distributed as follows were analyzed: 11 females and 11 males originated from the isolines (total of 22); 3 females and 7 males collected from the field in 2024; 20 females collected from the field in 2026 randomly. Besides, 20 *Chartocerus* individuals (10 females and 10 males) were analyzed. To prevent artefactual detection of viruses in parasitoids and hyperparasitoids due to the phoretic presence of viral particles derived from their host, each individual was rinsed with PBS to remove any residual biological material from the host body. Total RNAs were extracted individually as described above. Following, cDNA synthesis was performed on using Prime Script RT reagent kit (Takara Bio Inc., Japan). RT-qPCR was performed in a QuantStudio 3 Real-Time PCR System (Applied Biosystems, Thermo Fisher Scientific, Waltham, MA, USA) using hot firepol EvaGreen qPCR mix Plus (ROX) from Solis BioDyne (Estonia). Specific primers for all viruses characterized, as well as housekeeping genes for each species were used for viral detection and quantification (Table S2). The housekeeping gene of each insect species was amplified as an endogenous control of RNA concentration. Prior to use, the efficiency and specificity of each primer pair (viral and housekeeping) were evaluated. A sample was considered positive for a given virus when it showed a single melting peak matching that of the specific viral amplicon and a Ct value below 35; samples not meeting both criteria were considered negative (not detected). The relative viral abundance represents the abundance of viral molecules in relation to the abundance of the housekeeping gene and was calculated by comparing viral and housekeeping Ct values after adjusting for primer efficiency (*34*). Relative abundances were log10-transformed, and non-detected samples were assigned a value of −5.

The viruses ADiV1 and PsAmV1 were selected to study interspecies viral transmission based on their presence in both *Anagyrus* species. RNA sequencing samples from *P. citri, D. aberiae, A. aberiae,* and three *A. vladimiri* strains were mapped against the Anagyrus dicistrovirus 1 (ADiV1) and Pseudococcidae alphamesonivirus 1 (PsAmV1) reference genomes using BWA-MEM (*35*). Parasitoids from strains A and B were obtained from two commercial biological control companies in Spain whereas data for strain C, a population from the Middle East, were obtained from the publicly available NCBI BioProject PRJNA641546. Alignments were filtered to retain reads with a minimum mapping quality (Q > 20), converted to BAM format, sorted, and indexed using SAMtools (*36*). For each sample, variants were called against the virus reference genome using BCFtools (mpileup + call) (*37*). Variant calls for the target virus were extracted, and consensus sequences were generated with bcftools consensus. Per-base coverage was calculated with SAMtools depth, and coverage metrics (mean depth, breadth ≥1× and ≥10×) were computed. Low-confidence positions with depth below a defined threshold (< 3) were masked with N, and only these depth-masked consensus sequences were used for downstream analyses. Coding sequences (CDS) were extracted and concatenated from each consensus sequence and aligned using TranslatorX (http://translatorx.co.uk) to produce codon-aware nucleotide alignments. Maximum-likelihood phylogenies with IQ-TREE2 (v2.x) (*32*), applying ModelFinder to select the best-fit substitution model automatically (-m MFP) and assessing branch support with 1000 ultrafast bootstrap replicates (-B 1000).

### Cohabitation assay

To assess whether cohabitation could serve as a route of viral transmission between *A. vladimiri* and *A. aberiae,* unmated 2-day-old *A. vladimiri* (obtained from a commercial line) and *A. aberiae* (from line 17/19UV) were used. Each female of an *Anagyrus* species was placed in a 2 mL glass vial for 48h with a male of the other *Anagyrus* species allowing cohabitation. Each vial contained 70:30 (v/v) honey–water droplets and was closed with a cotton wool ball. After 48h, individuals were exchanged between pairs placing each female in a 2 mL glass vial with a male of their respective species that was previously cohabitating 48h with a female of the other species and were allowed to mate for 24h. Following, males were frozen at –20°C and females were introduced in a Petri dish with a mesh-covered lid to allow ventilation. In each Petri dish, 10 mealybugs suitable for parasitism (*P. citri* for *A. vladimiri* and *D. aberiae* for *A. aberiae*) were offered. Twenty-four hours later, female parasitoids were recovered and frozen at –20°C. Once their offspring emerged, individuals were sexed, recorded and frozen at –20°C. All parental and offspring individuals (G1) were analyzed for viral detection as described above.

### Multiparasitism assay

To assess the role of multiparasitism (when a single host individual is parasitized by two or more parasitoid species) as a potential mechanism for viral transmission between parasitoids, we designed an assay in which a mated *A. aberiae* female was allowed to oviposit a suitable parasitable stage of *P. citri* followed by a multiparasitism by *A. vladimiri*. *Anagyrus aberiae* specimens belong to mass rearing and *A. vladimiri* was provided by a biological control company. The mealybug was settled on a green bean pod and placed under the binocular. By visual observations, when *P. citri* was parasitized, *A. aberiae* was collected and frozen at –20 °C and the parasitized mealybug individual was moved to a ventilated Petri dish and incubated at 25 °C, 70%RH and 14:10 L:D regime. After 24h, the parasitized *P. citri* was offered to a mated *A. vladimiri* female. By visual observations under the binocular, when *P. citri* was parasitized*, A. vladimiri* was collected and frozen at –20 °C. Sixteen replicates were performed and upon emergence, each parasitoid (G_0_) was taxonomically identified.

To assess if the potential viral acquisition was vertically transmitted, *A. vladimiri* females from the G_0_ were individually paired for 24h with one conspecific male in glass vials with 70:30 (v/v) honey–water droplets. Subsequently, each female was released in a ventilated Petri dish with 20 mealybugs suitable for parasitism and allowed to parasitize for 24h. Afterwards, each female was individualized, frozen and screened for RNA viruses. Viral presence and abundance were assessed as described above. The immediate descendance (G1) of *Anagyrus* females that acquired a virus from the other *Anagyrus* species were selected, mated and allowed to parasitize the corresponding host for subsequent generations. Viral presence and abundance were assessed as described above.

### Impact of recent viral spillover event on the new host performance

To assess whether the acquisition of the Anagyrus vladimiri reovirus (AnvRV) affects the performance of its new host, we used an *A. aberiae* laboratory population (SelUV) in which AnvRV had been introduced experimentally through cohabitation with *A. vladimiri*. The virus became established in this population but did not reach fixation, and its prevalence varied among individuals. The effect of the virus was therefore assessed by direct comparison between AnvRV-positive and AnvRV-free females from the same cohort. The assay was conducted with a total of 155 mated, 7-day-old females. Each female was introduced individually into a 6 cm diameter ventilated Petri dish containing four females and four third-instar *D. aberiae* nymphs on a small piece of lemon and allowed to oviposit for 24 h. Females were then collected, frozen individually at –20 °C and later screened for AnvRV presence and relative abundance. Mealybugs were kept at 25 °C, 70%RH and 14:10 L:D regime until emergence of the offspring. Once offspring emerged, the number of parasitoids per female, sex ratio of the offspring and mealybug mortality were recorded. In parallel, 15 control Petri dishes were set up under identical conditions but without releasing a parasitoid female. Parasitoid emergence was analyzed with a generalized linear model (GLM) with a quasi-Poisson error distribution and log link function. Offspring sex ratio (females out of total emerged; plates with no emergence were excluded) and mealybug mortality (dead out of mealybugs offered) were analyzed as proportions with GLMs with a quasi-binomial error distribution and logit link function. Data dispersion was assessed by calculating the ratio of the sum of squared Pearson residuals to the residual degrees of freedom; because this ratio exceeded unity (φ = 1.19 for emergence and 2.27 for sex ratio), quasi-likelihood models were used so that standard errors accounted for among-female variation. Analyses were performed in R v4.3.3.

## Results

### Rich viral diversity in the *Anagyrus*-pseudococcid complex

Fifteen coding-complete different RNA viruses were identified and characterized across the two mealybug species (*D. aberiae*, *P. citri*) and their parasitoids (*A. aberiae* and *A. vladimiri*), thirteen of them being newly described (Table S3). Based on the RdRp phylogeny and their genomic structure, they were assigned to eleven viral families (Fig. S1, Table S3-S4).

Positive-sense single-stranded RNA viruses (+ssRNA) were the most abundant, comprising ten species across seven families: *Dicistroviridae* (n = 3), *Mesoniviridae*, *Flaviviridae*, *Picornaviridae*, *Iflaviridae*, *Virgaviridae* and *Tombusviridae* (n = 1 each) (Fig. 1B, Table S3). One additional virus is an unclassified member of the order *Nidovirales*. Three negative-sense single-strand RNA viruses (-ssRNA) belonged to two different families: *Phasmaviridae* (n=2) and *Rhabdoviridae* (n=1) (Fig. 1B, Table S3). Additionally, two double-strand RNA viruses (dsRNA) were identified, one assigned to the *Totiviridae* family and one to the *Sedoreoviridae* family (Fig. 1B, Table S3).

### Multiple viruses are shared across related species and two trophic levels

Our data reveal that some viruses can be found in multiple hosts: in phylogenetically related species and across two trophic levels (Fig. 1B). The most remarkable case occurs with the Pseudococcidae alphamesonivirus 1 (PsAmV1) which was independently detected in all four species. Furthermore, Delottococcus aberiae dicistrovirus 2 (DaDiV2) and Planococcus citri dicistrovirus 1 (PcDiV1) were each detected in three species, the two parasitoids and *D. aberiae* or *P. citri,* respectively. Other viral species were shared among more closely related hosts at the same trophic level: Planococcus citri flavivirus 1 (PcFlV1) and Pseudococcidae virgavirus 1 (PsViV1) were found exclusively in the two mealybug species, whereas ADiV1 was restricted to the two parasitoid species. The remaining nine viruses were detected each in a single host species.

Analysis of the viral abundance in parasitized mealybugs showed that each *Anagyrus* parasitoid transmitted a distinct viral set through oviposition, regardless of the mealybug host (Fig. S2). AaPhV1 (specific to *A. aberiae*) was detected at high abundance after parasitizing both mealybugs, even when the parasitoid egg had been removed at 36 h, indicating that the viral reads originate mainly from the mealybug tissues (Fig. S2A and B). The same pattern was observed for the AvPhV1, AvTV1, and AnvRV (specific to *A. vladimiri*) when *A. vladimiri* oviposits in either permissive (Fig. S2B) or non-permissive hosts (Fig. S2A). Transmission through oviposition is virus-specific since ADiV1 (specific to the parasitoid) was undetected in the host following oviposition (Fig. S2). These results showed the transference of viruses through parasitization, even in non-permissive hosts, indicating that shared space may contribute to the viral flow between the native and the exotic chains.

The genotypes of PsAmV1 recovered from the four analyzed species were over 99.9% identical at the nucleotide level, indicating that the same viral population circulates across native and invasive species and across both trophic levels. Sequence identity among genotypes of the other shared viruses was lower but still above 97.5%. Phylogenetic reconstruction of PsAmV1 genotypes revealed that PsAmV1 clusters according to their geographical origin rather than by host species: samples derived from the Spanish populations analyzed here (*A. vladimiri* strain A, *A. aberiae, D. aberiae* and *P. citri*) were more closely related to each other than to the PsAmV1 derived from *A. vladimiri* strain collected in the Middle East (PsAmV1 strain C) (Fig. 1C). Phylogenetic analysis of ADiV1, a virus exclusively found in the two *Anagyrus* species, showed the same phylogeographic structure (Fig. 1C). Together, these patterns suggest virus exchange among hosts within the Spanish agroecosystem, rather than independent introductions alongside each host lineage.

### Viral flow follows the *A. aberiae* introduction and extends to the hyperparasitoids

To determine whether viral flow followed the introduction of *A. aberiae,* we compared the viromes of parasitoid individuals from different origins: South African-originated isofemale lines (n=22), maintained independently since importation and never exposed to *A. vladimiri* or its host *P. citri*; and field-collected individuals from 2024 (n=10), and from 2026 (n=20). Only four viruses (AaPhV1, PsAmV1, AvNiLV1, and ADiV1) were detected in the isolines, whereas all nine viruses were detected among field-collected individuals, five of which were absent from the imported lines (Fig. 2A). Moreover, the viral repertoire of field populations expanded over time: none of the viruses restricted to *A. vladimiri* was detected in 2024, whereas all three (AvPhV1, AnvRV and AvTV1) were present in 2026. These results provide direct evidence of a recent, and most likely ongoing, viral flow between the native and the introduced parasitoid.

**Figure 2.**
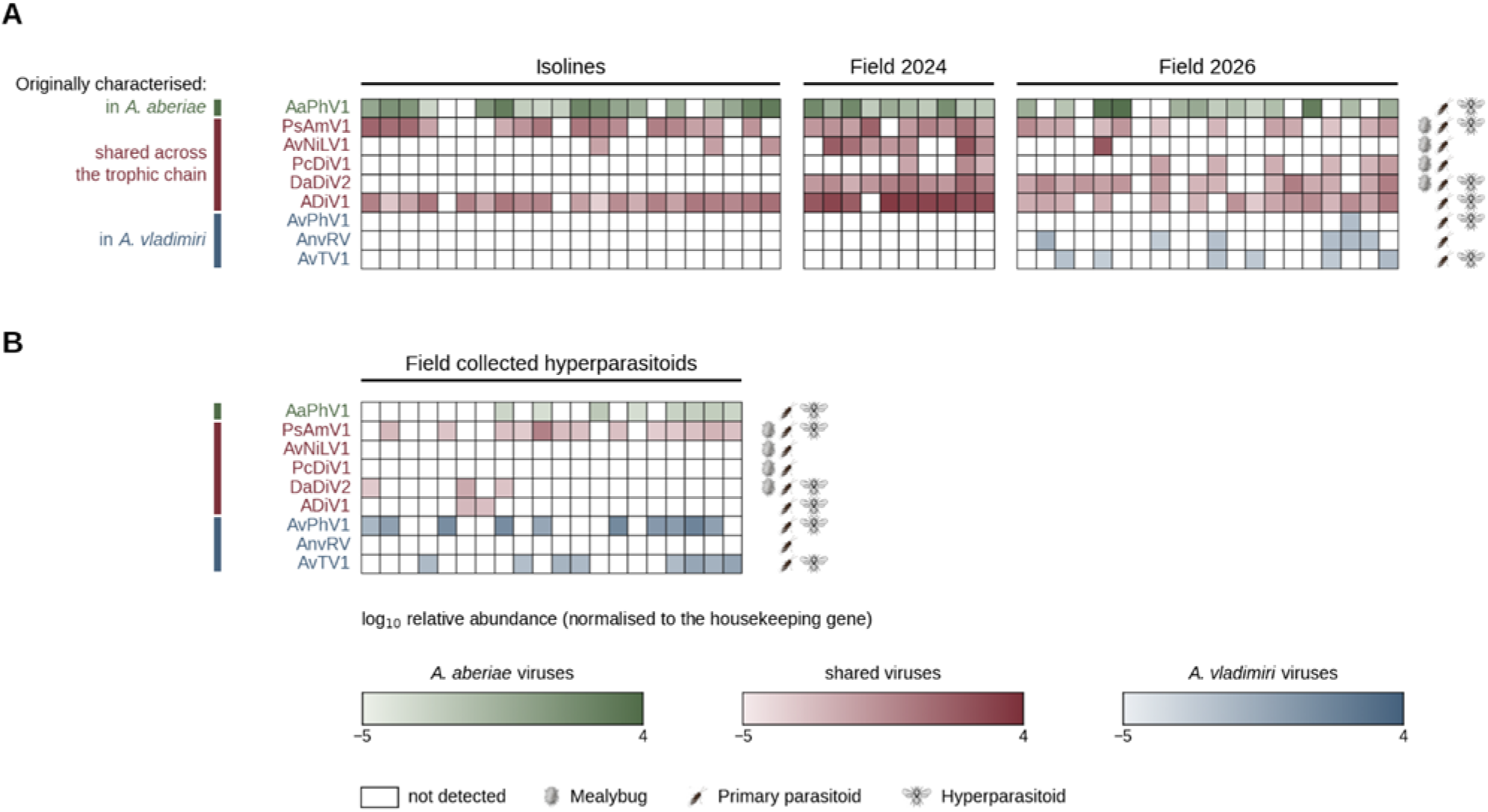
Viral flow across three trophic levels. **(A)** Heatmaps representing the relative abundance of viruses detected in *A. aberiae* isofemale lines and field-collected *A. aberiae* from 2024 and 2026. **(B)** Heatmap representing the relative abundance of viruses detected in field-collected hyperparasitoids (*Chartocerus* sp.). Each column represents one individual. Cell color intensity indicates log₁₀ relative abundance, quantified by qPCR and normalized to the housekeeping gene. White cells indicate that the virus was not detected. Viruses are grouped and colored according to where they were originally characterized: in *A. aberiae* (green), shared across the trophic chain (red), or in *A. vladimiri* (blue). Icons to the right of each row indicate the trophic level (TL) in which each virus has been detected: mealybug (2^nd^), primary parasitoid (3^rd^) and hyperparasitoid (4^th^).

We next asked whether viral flow extended to other members of the native trophic chain. We therefore screened field-collected individuals of the generalist hyperparasitoid *Chartocerus* sp. (n=20) from citrus orchards where *A. aberiae* was well established. Viruses shared across the trophic network (PsAmV1, ADiV1, and DaDiV2) and viruses restricted to *A. vladimiri* (AvPhV1 and AvTV1) were detected across individuals (Fig. 2B). Notably, the only virus exclusively associated with *A. aberiae* (AaPhV1) was also found (8 of 20 individuals), showing that a virus co-introduced with the exotic parasitoid reached the top of the native trophic network.

### Cohabitation and multiparasitism as putative routes of viral flow

To identify possible mechanisms underlying the observed viral flow, we evaluated cohabitation and multiparasitism as two potential transmission routes.

To mimic the cohabitation scenario, individuals from one species were allowed to cohabit for 48h with opposite-sex individuals of the other species and subsequently mated with conspecific partners. Both parental pairs and their offspring were analyzed (Fig. 3A). Viral transmission occurred in both directions. AvPhV1 (specific for *A. vladimiri*) was detected in 6 of 14 (43 %) of *A. aberiae* females and 6 of 15 (40 %) males. AnvRV (specific for *A. vladimiri*) was detected in 3 of 14 (21%) females and 6 of 15 (40%) males. Conversely, AaPhV1 (specific for *A. aberiae*) was detected in 8 of 15 (53%) of *A. vladimiri* females and 12 of 15 (80%) of males. Heterospecific viruses were further recovered from the G1 offspring of both *A. aberiae* and *A. vladimiri* founder females (Fig. 3A). AaPhV1 in 5 of 21 (24%) G1 individuals from *A. vladimiri* mothers, and AnvRV and AvTV1 in 6 of 19 (32%) each from *A. aberiae* mothers, although AvPhV1 was not recovered from any of the latter. These results show that viruses acquired through cohabitation are passed to the next generation and confirm that the mere co-occurrence of the two species is sufficient to drive viral flow in both directions.

**Figure 3.**
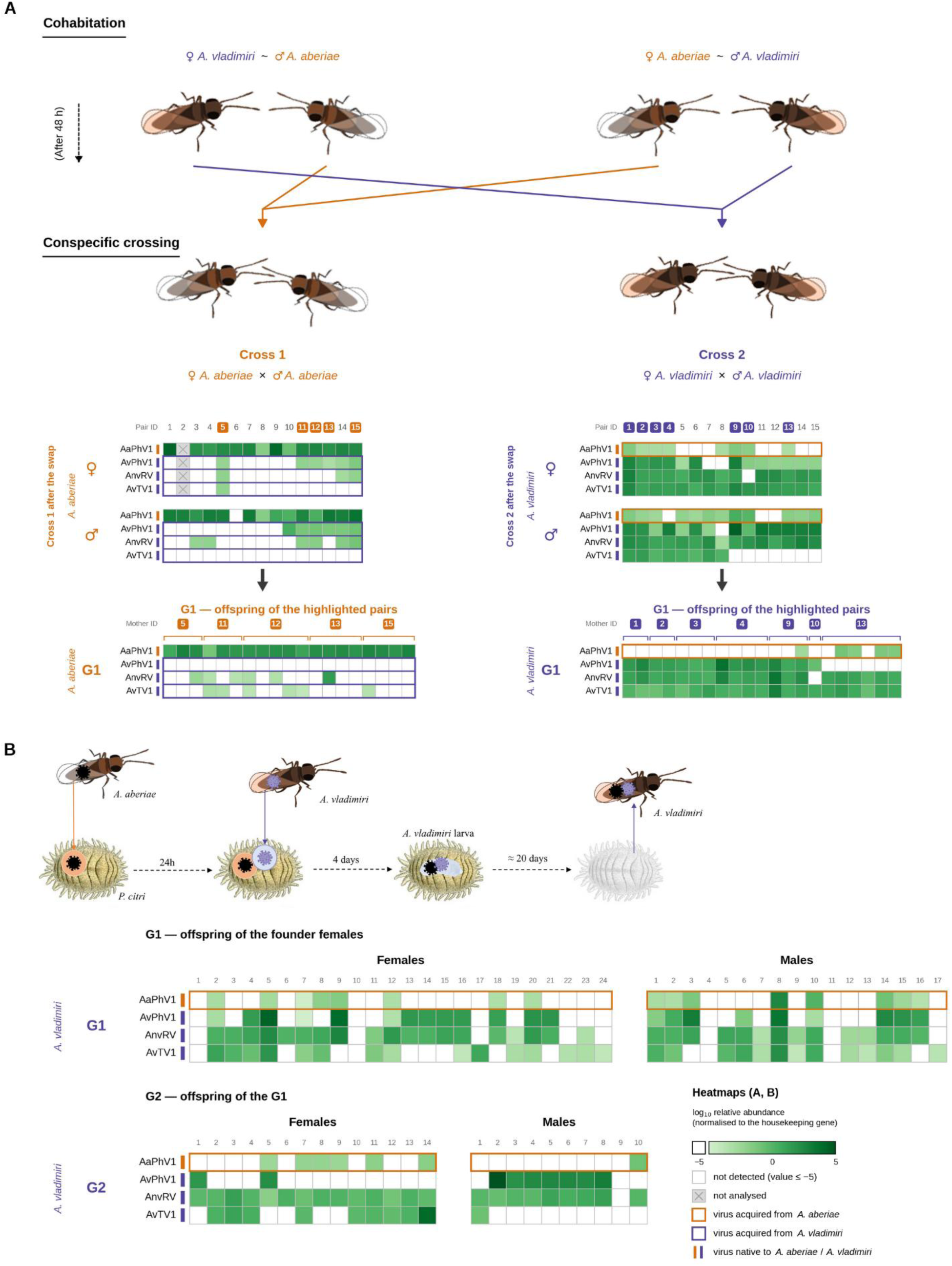
Cohabitation and multiparasitism as routes of viral flow between species. **(A)** Cohabitation assay. Females and males of *A. aberiae* and *A. vladimiri* were held as heterospecific pairs for 48 hours and then mated with conspecific partners, yielding Cross 1 (*A. aberiae*) and Cross 2 (*A. vladimiri*). Heatmaps represent the prevalence and abundance of transmitted viruses between cohabiting *A. aberiae* and *A. vladimiri* individuals and their immediate progeny (G1). Cell color intensity indicates log₁₀ relative abundance, quantified by qPCR and normalized to the housekeeping gene. Highlighted pair IDs indicate the females whose offspring were screened. G1 individuals are grouped by mothers. **(B)** Multiparasitism assay. An *A. aberiae* female oviposited in a *Planococcus citri* nymph, an *A. vladimiri* female oviposited in the same host 24h later, and the emerging *A. vladimiri* was screened. Heatmaps show the viruses detected in the G1 and G2 descendants of three *A. vladimiri* founder females that had acquired AaPhV1 through multiparasitism. Cell color intensity indicates log₁₀ relative abundance, quantified by qPCR and normalized to the housekeeping gene.

We next examined multiparasitism as a driver of viral flow. Here, a single host is sequentially parasitized by more than one parasitoid species. Because both *Anagyrus* species are solitary, only one larva completes development, and the surviving parasitoid may acquire viral particles while feeding on a host already containing the embryo of the competitor (Fig. 3B). *Planococcus citri* individuals parasitized by *A. aberiae* were offered to *A. vladimiri* females 24h later. Of the 16 replicates, five adults emerged, and three females were identified as *A. vladimiri* (males cannot be distinguished between the two species). All the three females carried AaPhV1, the *A. aberiae* specific virus. To determine whether the acquired virus was vertically transmitted rather than merely carried over from the host, we screened the descendants of these three founder females after mating with *A. vladimiri* males from the colony (Fig. 3B). AaPhV1 was detected in 8 of 24 (33%) G1 females and 8 of 17 (47%) males, and persisted in the second generation, being detected in 7 of 24 (29%) G2 descendants (Fig. 3B). Persistence across two generations indicates genuine infection of the new host rather than residual host material. Notably, individuals carrying AaPhV1 retained their resident *A. vladimiri* viruses (AvPhV1, AnvRV, and AvTV1), indicating that acquisition does not displace the established virome (Fig. 3B). Together these experiments show that AaPhV1, co-introduced with *A. aberiae,* readily spills over to the native *A. vladimiri*.

### Acquired virus enhances the performance of its new host

Having detected the recent acquisition of AnvRV by field populations of *A. aberiae* (Fig. 2A), we set out to determine whether viral flow across the trophic network had consequences for the recipient host. AnvRV was originally described in *A. vladimiri,* where it facilitates egg hatching by reducing its encapsulation in *P. citri* (*38*). We, therefore, established an *A. aberiae* laboratory colony in which AnvRV had been introduced experimentally through cohabitation with *A. vladimiri*. Because the virus did not reach fixation in this colony, the effect of the virus was obtained by direct comparison between AnvRV-positive and AnvRV-free females from the same cohort.

Females were allowed to parasitize their natural host, *D. aberiae,* and offspring emergence was recorded. AnvRV-positive females produced more offspring than virus-free females (4.55 ± 0.40 *vs* 3.49 ± 0.19 emerged parasitoids per female, a 30% increase; quasi-Poisson GLM, *P* = 0.016) (Fig. 4A), without affecting the sex ratio (0.73 ± 0.06 vs 0.68 ± 0.03 female proportion; quasi-binomial GLM, *P* = 0.29) (Fig. 4B) or host mortality (0.65 ± 0.14 vs 0.80 ± 0.08 dead *D. aberiae* individuals per plate; quasi-binomial GLM, *P* = 0.32) (Fig. 4C). No mealybug mortality was recorded in the controls. Therefore, AnvRV retains its beneficial effect in the new host following spillover, without a detectable cost to host exploitation or to offspring sex allocation, reflecting that viral flow across a trophic network shapes the relations among the members and is relevant for the successful introduction of novel biological control agents.

**Figure 4.**
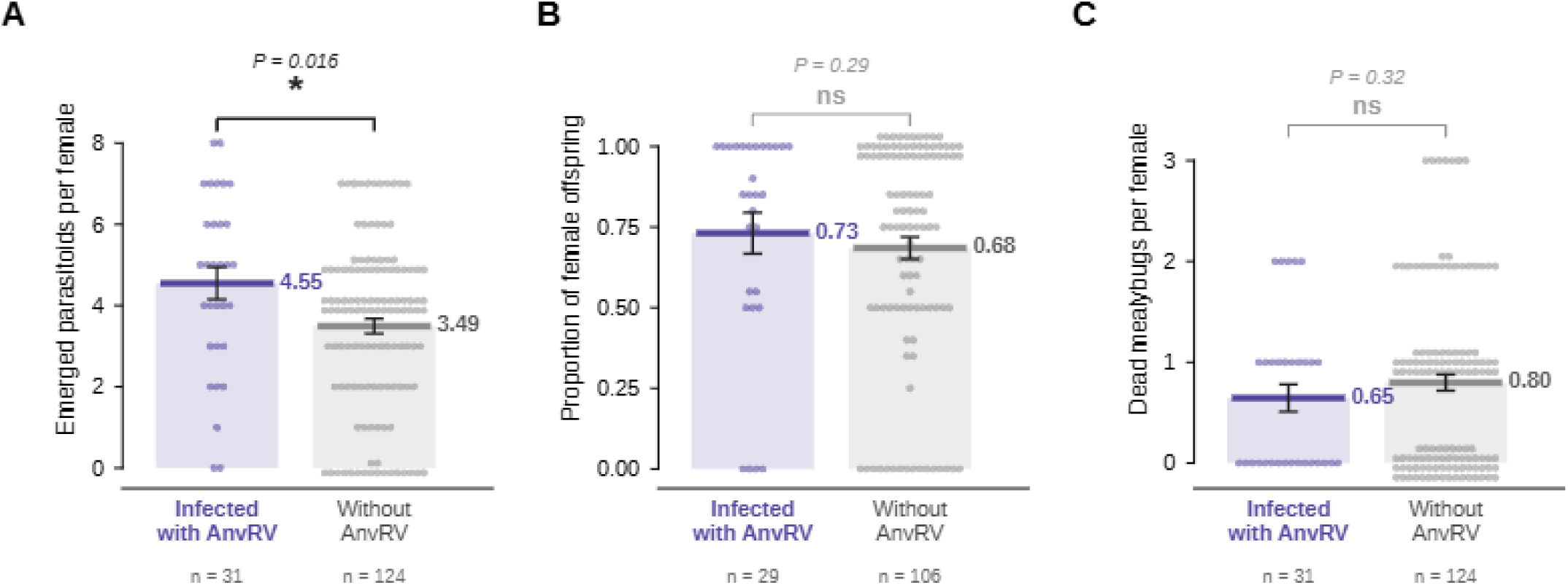
Acquisition of AnvRV enhances parasitism in *A. aberiae*. **(A)** Parasitoid offspring emerging per female. **(B)** Proportion of female offspring (females producing no offspring are excluded). **(C)** Dead *D. aberiae* per female. Bars show group means ± standard error, and each point represents one female. \**P* < 0.05; ns = not significant. Sample sizes are given below each bar.

## Discussion

Species introductions, whether accidental or deliberate, have traditionally been framed in terms of competition, predation and adaptation to novel habitats (*12*, *39*). A largely overlooked consequence is that every introduction also delivers a community of associated microorganisms into the recipient environment, including cryptic viruses whose fate has remained unexplored. Here, we show that when previously isolated species are brought into contact, their associated viruses flow bidirectionally between exotic and native trophic chains. This viral flow spanned three consecutive trophic levels, from mealybugs to the hyperparasitoids at the fourth trophic level, and had consequences, in our case, conferring a measurable advantage on the newly infected host.

Across the four species examined, we identified fifteen RNA viruses, some of them shared among two or more species. Although for most of them we cannot reconstruct the origin or the direction of their acquisition, three independent lines of evidence of viral flow among species converge for a subset of viruses: (i) the genotypes of the most widely distributed viruses cluster by geographical origin rather than by host species, a pattern expected from local and recent exchange but not from viruses carried in by each population; (ii) the South African isofemale lines provide a reference against which field populations show the acquisition of viruses previously restricted to the native chain (AvPhV1, AnvRV, and AvTV1); and (iii) cohabitation and multiparasitism, two interactions that occur naturally, were sufficient for viruses to be transferred between species and maintained transgenerationally. Together, these observations capture viral flow as an ongoing process with the potential to reshape trophic network dynamics.

When an exotic trophic chain co-occurs with a native one, the previously existing ecological barriers disappear, making host shifting feasible (*40*). Our cohabitation assay confirms that close contact between the two *Anagyrus* species is enough for viruses to transmit in both directions, and vertically within the new host population. Multiparasitism, which occurs naturally when suitable hosts are scarce (*41*), provides an alternative route. Although the two *Anagyrus* species seldom complete development on each other’s host, they still oviposit in them (*42*) and our cross-oviposition analysis shows that they inject their associated viruses when they do. A multiparasitized mealybug, therefore, carries viruses of both species and since only one larva survives in solitary parasitoids (*43*), the survivor may acquire them through direct contact and/or feeding. Together, these two routes indicate that viral exchange between chains depends mainly on their co-occurrence.

An additional route of viral flow suggested by our data involves participation of a higher trophic level. *Chartocerus* sp. is a generalist hyperparasitoid that develops on primary parasitoids of both systems and on related species (*44*). The field-collected individuals we analyzed carried viruses from both trophic chains, including AaPhV1, which is specific to *A. aberiae* and was absent from this network before its release. We did not test whether these viruses replicate and transmit within the hyperparasitoid, although both possibilities could mediate viral mechanical transmission. Parasitoids probe more hosts than they successfully develop on, and failed attempts may transfer virions between them (*45*), regardless of whether they replicate. *Chartocerus* sp. also attacks closely related parasitoids in other trophic chains (*44*), which may contribute to a broader viral flow between food chains that are otherwise independent.

Viral transfer can produce measurable phenotypic changes in the new host and its interactions. AnvRV was initially restricted to *A. vladimiri*, being absent from the two mealybugs, the hyperparasitoid, and both the South African isolines and the 2024 field collections of *A. aberiae*. Two years later, it appeared in *A. aberiae* field collected individuals and our experiments have shown that it is prone to be acquired by *A. aberiae* through both cohabitation and multiparasitism. This virus is of particular interest as it reduces *A. vladimiri* egg encapsulation in *P. citri*, enhancing parasitism success (*38*). Once acquired by *A. aberiae*, AnvRV enhanced parasitoid success by increasing the adult emergence from *D. aberiae*, conferring a measurable advantage that might result in higher potential to control *D. aberiae* populations. Yet, newly established host-virus interactions might span across the mutualism-parasitism spectrum, and other acquisitions could affect their interactions differently.

Attention to viral spillover has so far focused on vertebrates, with predictions of around 4,000 cross-species viral transmission events among mammals alone by 2070 (*46*). Insects have mostly been studied as vectors of these pathogens, rather than as hosts of their own viral communities (*47*), leaving the viral diversity they introduce and its consequences across trophic levels unexamined. Deliberate introductions of natural enemies for pest control alone account for 2,384 insect species across 148 countries (*20*), likely co-introducing an enormous diversity of viruses. Our study supports the view that microorganisms, and viruses in particular, should be incorporated into invasion biology and biological control frameworks, as they are dynamic components of ecological communities capable of reshaping species interactions, ecosystem functioning and evolutionary trajectories following introductions.

## Credit authorship contribution statement

J.P-R: Conceptualization, experimental design, data collection, data analyses and writing – original draft. L.H-P: data analysis. R. G-G: Data collection, E. M-S: Data collection A.T: providing material, conceptualization. L.G: Conceptualization, experimental design, writing – review and editing; S.H: Conceptualization, experimental design, writing – review and editing.

## Acknowledgements

We would like to thank the State Insectary of Silla (Valencia, Spain), TRAGSA and biological control companies for insect supply. This research was supported by the Spanish Ministry of Science, Innovation and Universities under projects PID2021-124813OB-C33, PID2024-162646OA-I00 and PID2024-162058OB-C32, and the Ramón y Cajal grant (RYC2022-037286-I) MICIU/AEI/10.13039/501100011033. J. Pérez-Rodríguez was supported by the Spanish Juan de la Cierva postdoctoral program (FJC2021-046943-I), funded by MCIN/AEI/10.13039/501100011033.

## Supplementary material

**Figure S1.**
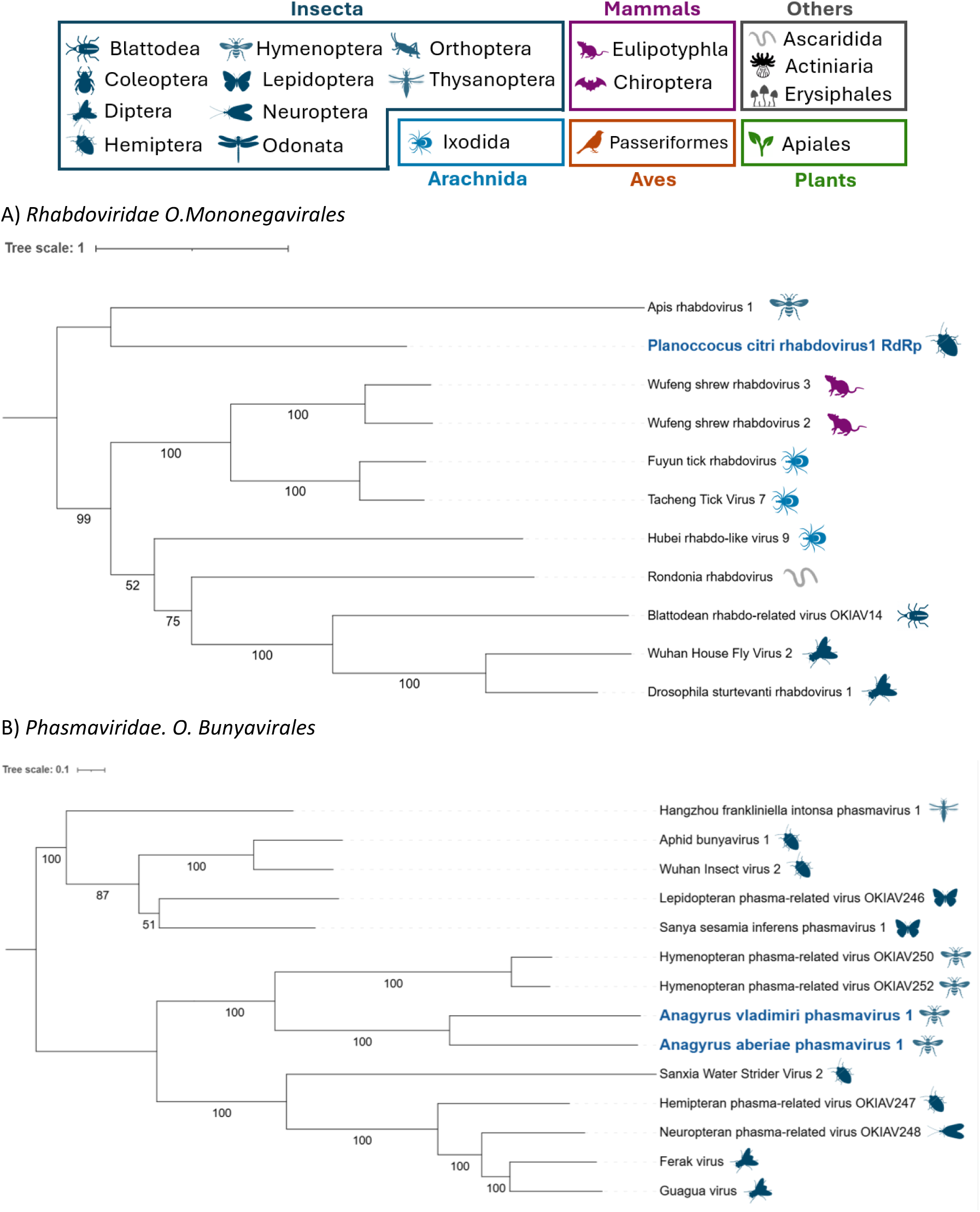
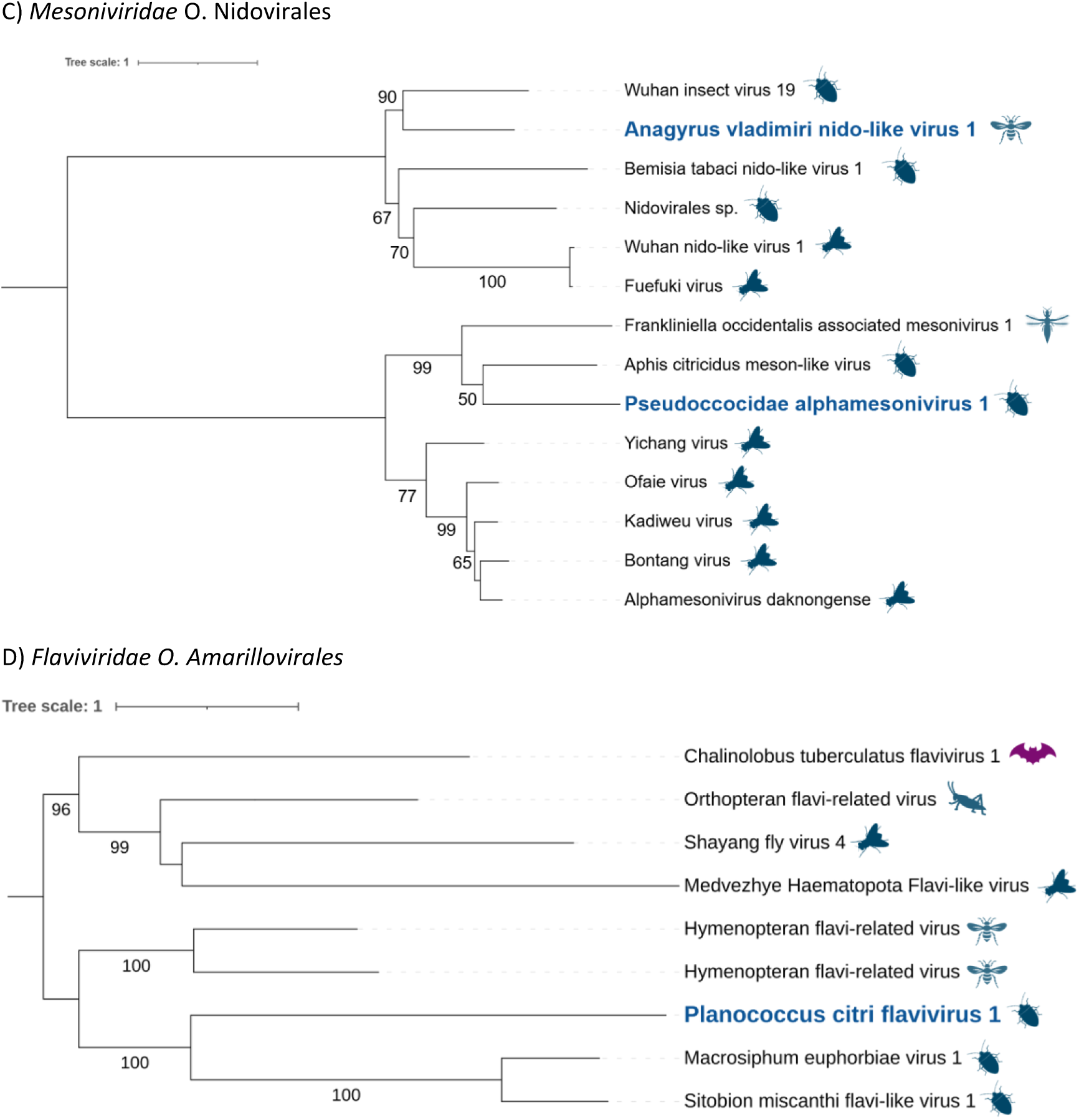
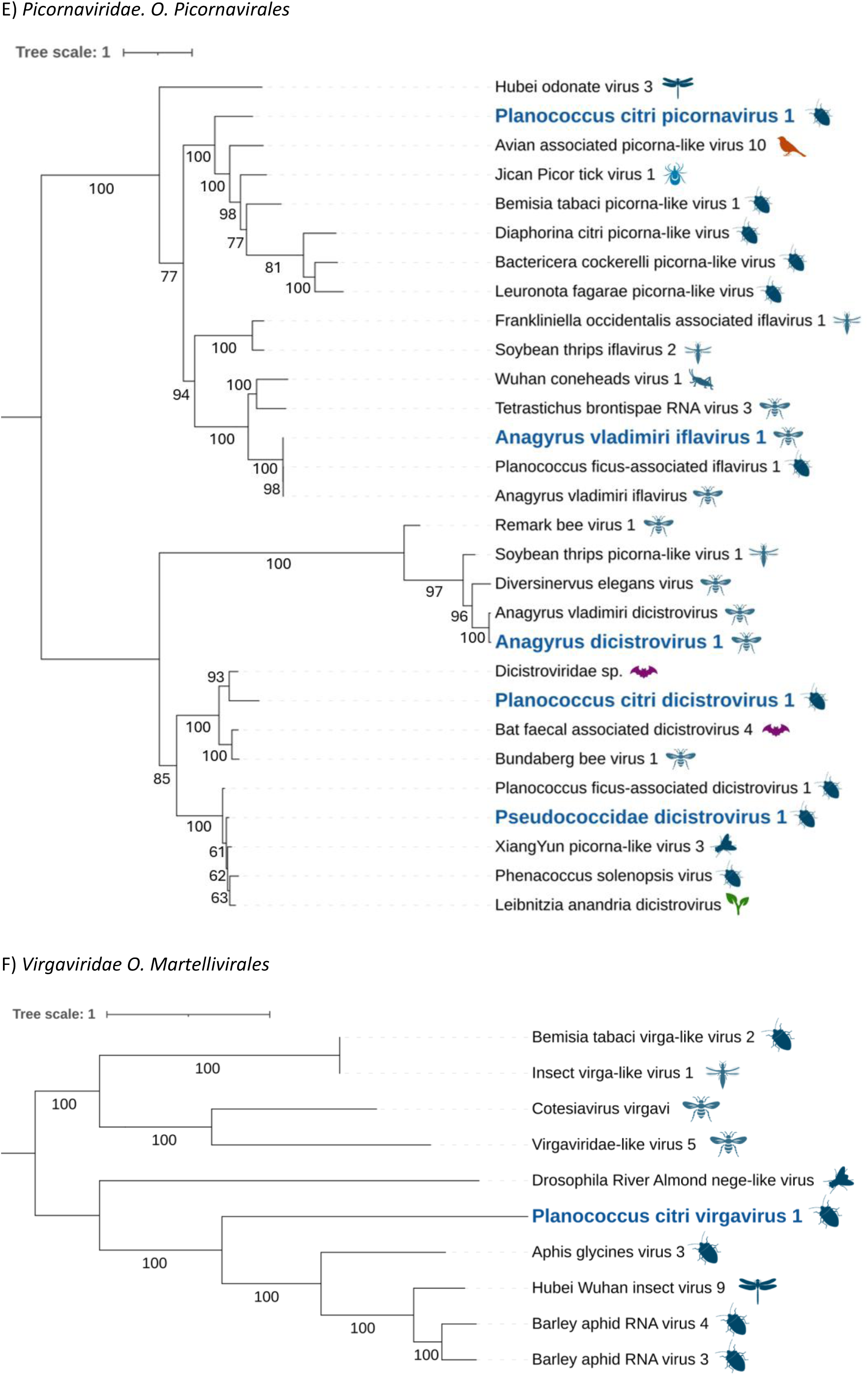
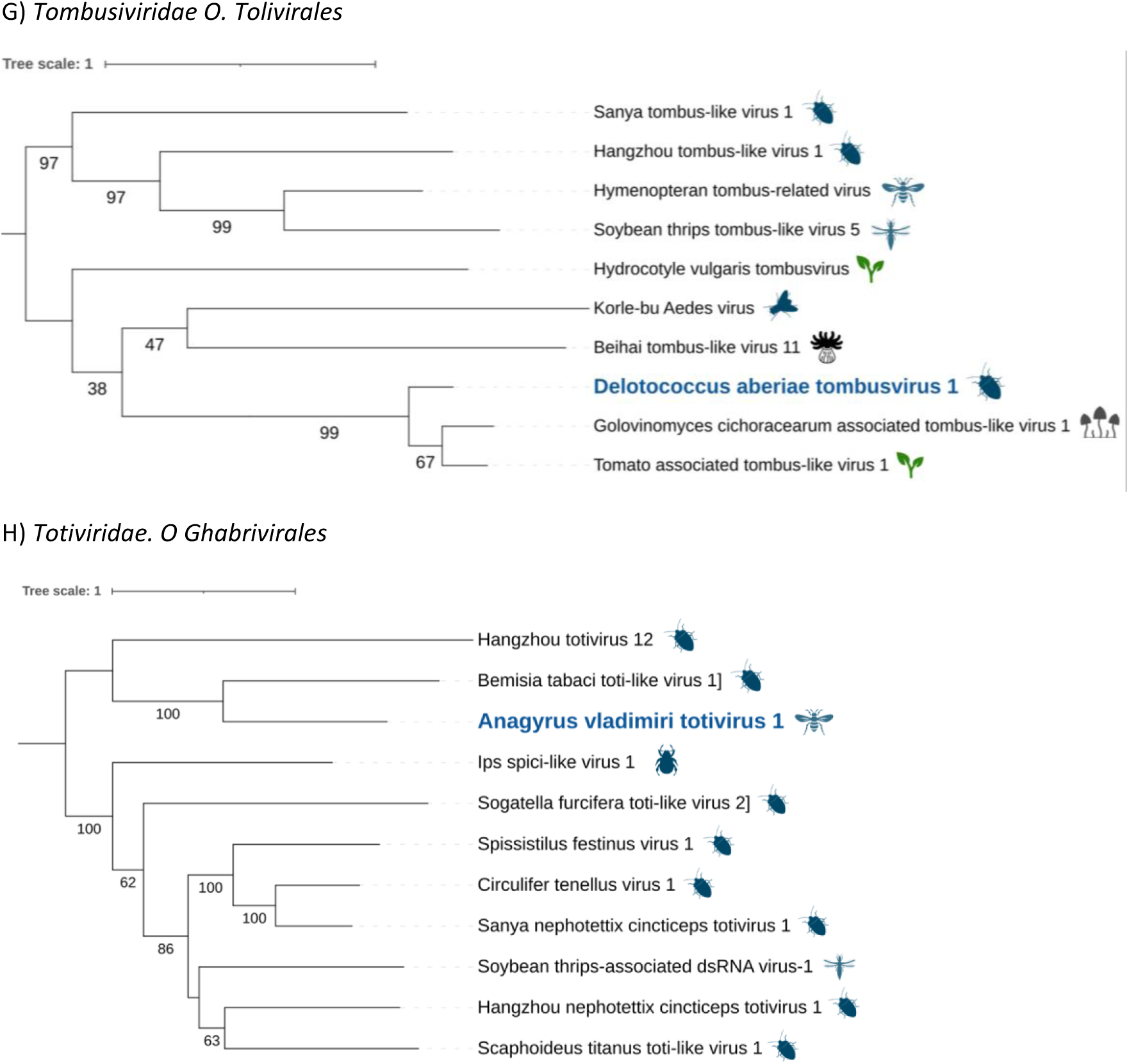
Phylogenetic classification of the newly described RNA viruses (in blue) in comparison to their closest viral relatives (in black). The closest viral relatives were detected using the viral RdRp region and blastp searches (Table S1). Phylogenetic inferences were calculated using IQtree2 with default settings and the consensus tree constructed from 1000 ultrafast bootstraps is shown. Symbols represent the host taxonomic order in which each of the virus was described (as specified in the legend on top of the figure).

**Figure S2.**
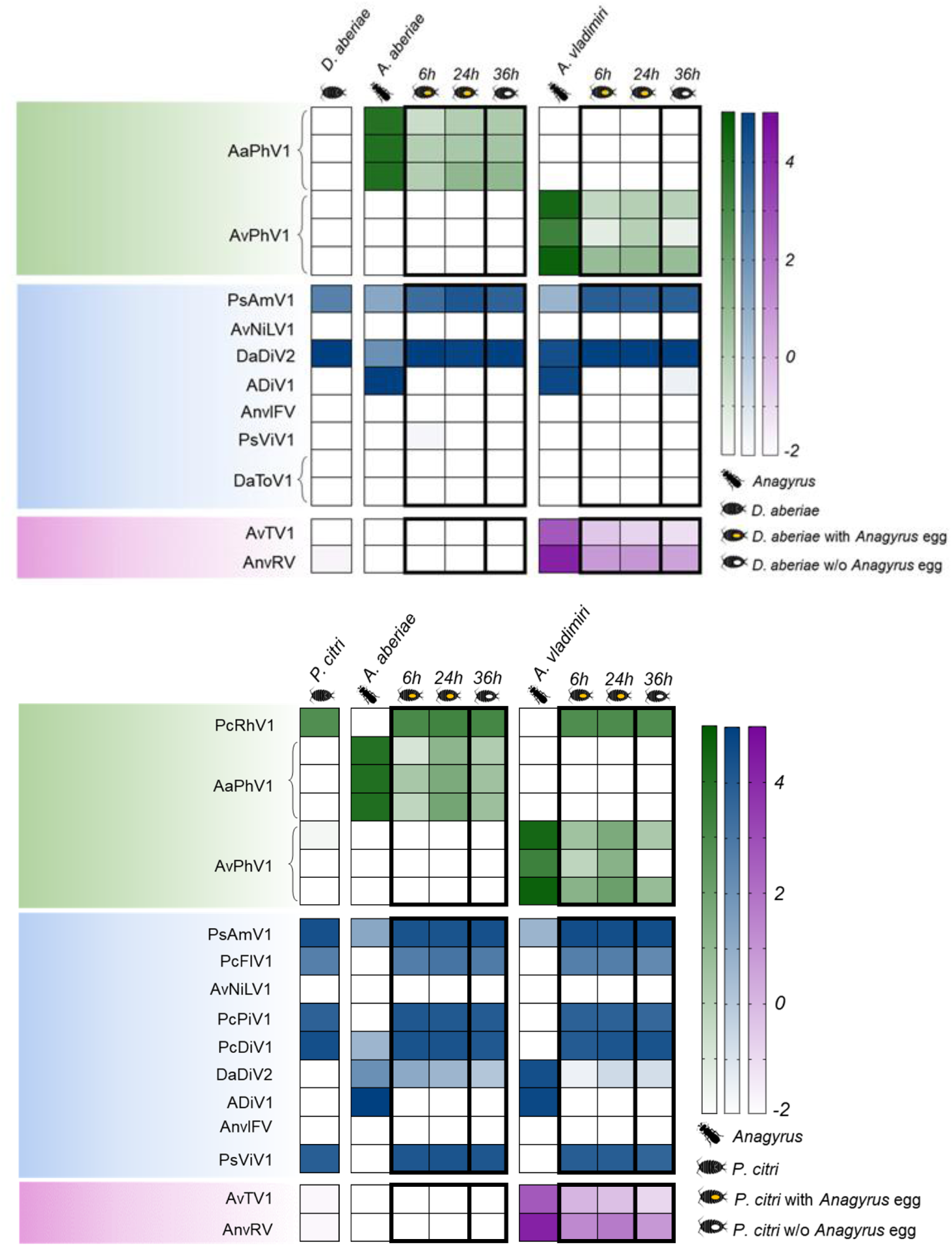
Heatmap representing the relative abundance of viruses detected when the host was. A) *Delottococcus aberiae* and B) *Planococcus citri.* Color intensity represents scaled relative abundance, from low (light) to high (dark). Each raw represents a virus and the columns represent the presence and abundance of viruses in the non-parasitized host, *A. aberiae* or *A. vladimiri* before oviposition and parasitized mealybugs 6h, 24h and 36h (when parasitoid egg was removed) post oviposition

**Table S1.** List of SRA accession numbers for each transcriptome sample.

| SRA number | Sample |
|---|---|
| SAMN45150978 | *P. citri* |
| SAMN45150979 | *P. citri* parasitised by *A. aberiae* 6h |
| SAMN45150980 | *P. citri* parasitised by *A. aberiae* 24h |
| SAMN45150981 | *P. citri* parasitised by *A. aberiae* 36h w/o egg |
| SAMN45150982 | *P. citri* parasitised by *A. vladimiri* 6h |
| SAMN45150983 | *P. citri* parasitised by *A. vladimiri* 24h |
| SAMN45150984 | *P. citri* parasitised by *A. vladimiri* 36h w/o egg |
| SAMN45150985 | *D. aberiae* |
| SAMN45150986 | *D. aberiae* parasitised by *A. aberiae* 6h |
| SAMN45150987 | *D. aberiae* parasitised by *A. aberiae* 24h |
| SAMN45150988 | *D. aberiae* parasitised by *A. aberiae* 36h w/o egg |
| SAMN45150989 | *D. aberiae* parasitised by *A. vladimiri* 6h |
| SAMN45150990 | *D. aberiae* parasitised by *A. vladimiri* 24h |
| SAMN45150991 | *D. aberiae* parasitised by *A. vladimiri* 36h w/o egg |
| SAMN45150992 | *A. aberiae* |
| SAMN45150993 | *A. vladimiri* |

**Table S2.**
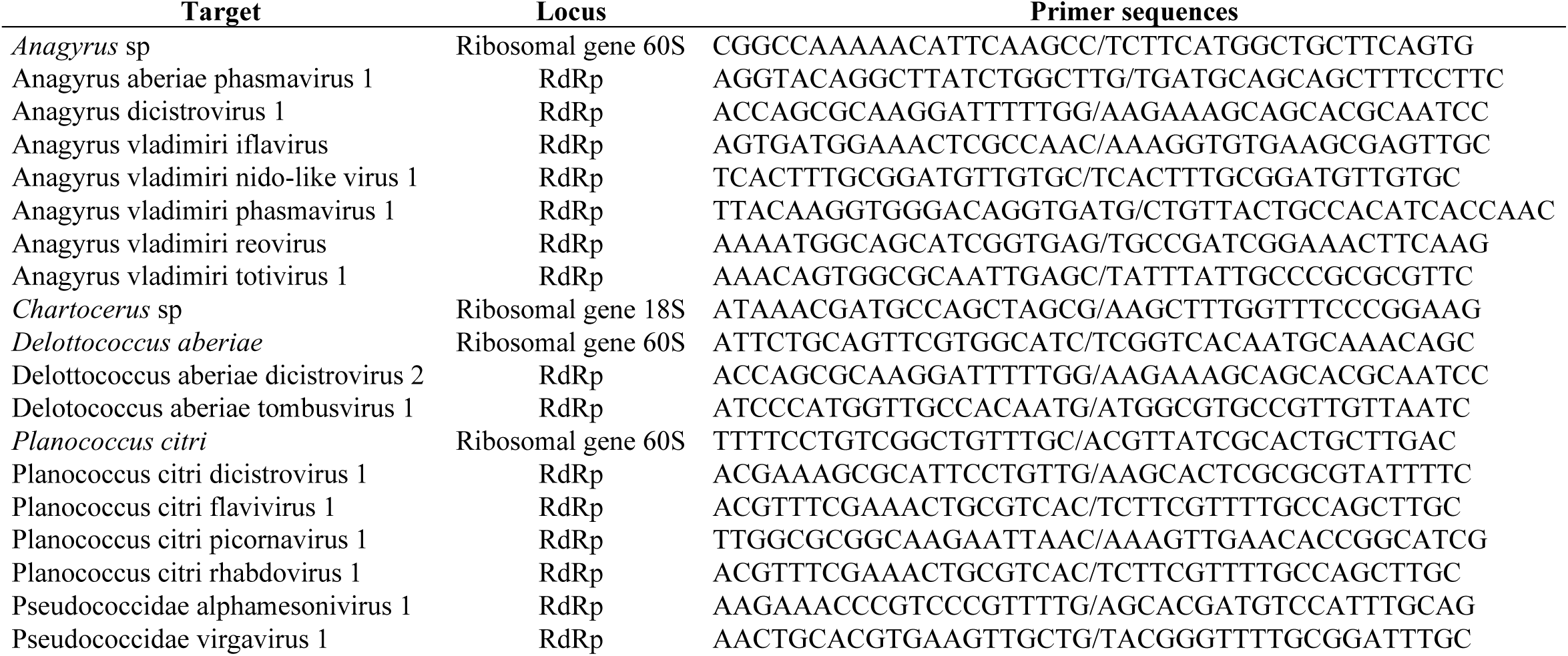
List of PCR primers used to detect and amplify the characterized RNA viruses and housekeeping genes.

**Table S3.**
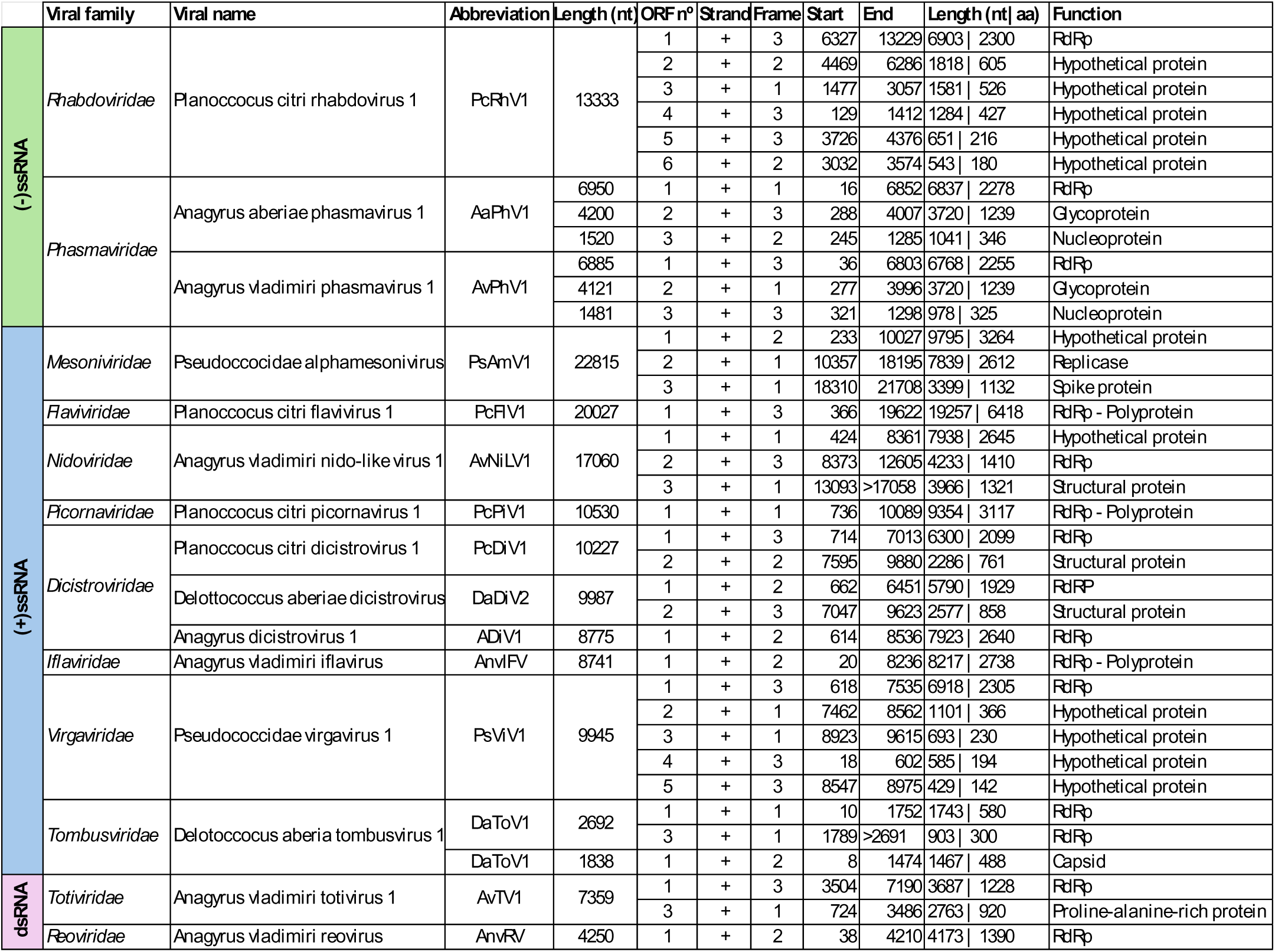
List of the 15 RNA viruses described in the *Anagyrus – Pseudoccocidae* complex clustered according to their genome type, viral family, genome length ORF number and function.

|  | Viral family | Viral name | Abbreviation | Length (nt) | ORF nº | Strand | Frame | Start | End | Length (nt\| aa) | Function |
|---|---|---|---|---|---|---|---|---|---|---|---|
| (-)ssRNA | *Rhabdoviridae* | Planoccocus citri rhabdovirus 1 | PcRhV1 | 13333 | 1 | + | 3 | 6327 | 13229 | 6903 \| 2300 | RdRp |
|  |  |  |  |  | 2 | + | 2 | 4469 | 6286 | 1818 \| 605 | Hypothetical protein |
|  |  |  |  |  | 3 | + | 1 | 1477 | 3057 | 1581 \| 526 | Hypothetical protein |
|  |  |  |  |  | 4 | + | 3 | 129 | 1412 | 1284 \| 427 | Hypothetical protein |
|  |  |  |  |  | 5 | + | 3 | 3726 | 4376 | 651 \| 216 | Hypothetical protein |
|  |  |  |  |  | 6 | + | 2 | 3032 | 3574 | 543 \| 180 | Hypothetical protein |
|  | *Phasmaviridae* | Anagyrus aberiae phasmavirus 1 | AaPhV1 | 6950 | 1 | + | 1 | 16 | 6852 | 6837 \| 2278 | RdRp |
|  |  |  |  | 4200 | 2 | + | 3 | 288 | 4007 | 3720 \| 1239 | Glycoprotein |
|  |  |  |  | 1520 | 3 | + | 2 | 245 | 1285 | 1041 \| 346 | Nucleoprotein |
|  |  | Anagyrus vladimiri phasmavirus 1 | AvPhV1 | 6885 | 1 | + | 3 | 36 | 6803 | 6768 \| 2255 | RdRp |
|  |  |  |  | 4121 | 2 | + | 1 | 277 | 3996 | 3720 \| 1239 | Glycoprotein |
|  |  |  |  | 1481 | 3 | + | 3 | 321 | 1298 | 978 \| 325 | Nucleoprotein |
| (+)ssRNA | *Mesoniviridae* | Pseudoccocidae alphamesonivirus | PsAmV1 | 22815 | 1 | + | 2 | 233 | 10027 | 9795 \| 3264 | Hypothetical protein |
|  |  |  |  |  | 2 | + | 1 | 10357 | 18195 | 7839 \| 2612 | Replicase |
|  |  |  |  |  | 3 | + | 1 | 18310 | 21708 | 3399 \| 1132 | Spike protein |
|  | *Flaviviridae* | Planoccocus citri flavivirus 1 | PcFlV1 | 20027 | 1 | + | 3 | 366 | 19622 | 19257 \| 6418 | RdRp - Polyprotein |
|  | *Nidoviridae* | Anagyrus vladimiri nido-like virus 1 | AvNiLV1 | 17060 | 1 | + | 1 | 424 | 8361 | 7938 \| 2645 | Hypothetical protein |
|  |  |  |  |  | 2 | + | 3 | 8373 | 12605 | 4233 \| 1410 | RdRp |
|  |  |  |  |  | 3 | + | 1 | 13093 | >17058 | 3966 \| 1321 | Structural protein |
|  | *Picornaviridae* | Planoccocus citri picornavirus 1 | PcPiV1 | 10530 | 1 | + | 1 | 736 | 10089 | 9354 \| 3117 | RdRp - Polyprotein |
|  | *Dicistroviridae* | Planoccocus citri dicistrovirus 1 | PcDiV1 | 10227 | 1 | + | 3 | 714 | 7013 | 6300 \| 2099 | RdRp |
|  |  |  |  |  | 2 | + | 2 | 7595 | 9880 | 2286 \| 761 | Structural protein |
|  |  | Delottococcus aberiae dicistrovirus | DaDiV2 | 9987 | 1 | + | 2 | 662 | 6451 | 5790 \| 1929 | RdRP |
|  |  |  |  |  | 2 | + | 3 | 7047 | 9623 | 2577 \| 858 | Structural protein |
|  |  | Anagyrus dicistrovirus 1 | ADiV1 | 8775 | 1 | + | 2 | 614 | 8536 | 7923 \| 2640 | RdRp |
|  | *Iflaviridae* | Anagyrus vladimiri iflavirus | AnvIFV | 8741 | 1 | + | 2 | 20 | 8236 | 8217 \| 2738 | RdRp - Polyprotein |
|  | *Virgaviridae* | Pseudococcidae virgavirus 1 | PsViV1 | 9945 | 1 | + | 3 | 618 | 7535 | 6918 \| 2305 | RdRp |
|  |  |  |  |  | 2 | + | 1 | 7462 | 8562 | 1101 \| 366 | Hypothetical protein |
|  |  |  |  |  | 3 | + | 1 | 8923 | 9615 | 693 \| 230 | Hypothetical protein |
|  |  |  |  |  | 4 | + | 3 | 18 | 602 | 585 \| 194 | Hypothetical protein |
|  |  |  |  |  | 5 | + | 3 | 8547 | 8975 | 429 \| 142 | Hypothetical protein |
|  | *Tombusviridae* | Delotoccocus aberia tombusvirus 1 | DaToV1 | 2692 | 1 | + | 1 | 10 | 1752 | 1743 \| 580 | RdRp |
|  |  |  |  |  | 3 | + | 1 | 1789 | >2691 | 903 \| 300 | RdRp |
|  |  |  | DaToV1 | 1838 | 1 | + | 2 | 8 | 1474 | 1467 \| 488 | Capsid |
| dsRNA | *Totiviridae* | Anagyrus vladimiri totivirus 1 | AvTV1 | 7359 | 1 | + | 3 | 3504 | 7190 | 3687 \| 1228 | RdRp |
|  |  |  |  |  | 3 | + | 1 | 724 | 3486 | 2763 \| 920 | Proline-alanine-rich protein |
|  | *Reoviridae* | Anagyrus vladimiri reovirus | AnvRV | 4250 | 1 | + | 2 | 38 | 4210 | 4173 \| 1390 | RdRp |

**Table S4.** Reference sequences used for the phylogenetic analyses shown in Fig. S1. Viral names and GenBank protein accession numbers of the closest relatives of each newly described RNA virus, identified by blastp searches of the RNA-dependent RNA polymerase (RdRp) region against the NCBI non-redundant protein database. Sequences are grouped by viral family, corresponding to panels (A) to (I) of Fig. S1.

| Viral name | Accession n<sup>o</sup><br>(protein) | Viral name | Accession n<sup>o</sup><br>(protein) |
|---|---|---|---|
| **(A) *Rhabdoviridae*** |  | Hymenopteran flavi-related virus | QTJ63569.1 |
| Tacheng Tick Virus 7 | YP_009304476.1 | Chalinolobus tuberculatus flavivirus 1 | WPA70758.1 |
| Fuyun tick rhabdovirus | UXL90877.1 | Medvezhye Haematopota Flavi-like virus | WQM60689.1 |
| Wufeng shrew rhabdovirus 2 | WPV62755.1 | Shayang fly virus 4 | YP_009179225.1 |
| Wufeng shrew rhabdovirus 3 | WPV62759.1 | Hymenopteran flavi-related virus | QTJ63570.1 |
| Hubei rhabdo-like virus 9 | YP_009337815.1 | **(E) *Picornaviridae*** |  |
| Blattodean rhabdo-related virus OKIAV14 | QMP82336.1 | Jinan Picor tick virus 1 | UYL95405.1 |
| Drosophila sturtevanti rhabdovirus 1 | AMK09264.1 | Bemisia tabaci picorna-like virus 1 | QWC36467.1 |
| Rondonia rhabdovirus | QHI42114.1 | Avian associated picorna-like virus 10 | USL85481.1 |
| Apis rhabdovirus 1 | WAK79924.1 | Leuronota fagarae picorna-like virus | UNG39521.1 |
| Wuhan House Fly Virus 2 | YP_009304985.1 | Bactericera cockerelli picorna-like virus | WWZ81655.1 |
| **(B) *Phasmaviridae*** |  | Diaphorina citri picorna-like virus | ALJ52073.1 |
| Hymenopteran phasma-related virus OKIAV252 | YP_010840603.1 | Soybean thrips iflavirus 2 | QQN90111.1 |
| Hymenopteran phasma-related virus OKIAV250 | YP_010840610.1 | Frankliniella occidentalis associated iflavirus 1 | QNM37810.1 |
| Neuropteran phasma-related virus OKIAV248 | YP_010840047.1 | Hubei odonate virus 3 | YP_009337289.1 |
| Ferak virus | AKN56913.1 | Dicistroviridae sp. | XCO49052.1 |
| Hemipteran phasma-related virus OKIAV247 | YP_010840003.1 | Bat faecal associated dicistrovirus 4 | UVF62134.1 |
| Lepidopteran phasma-related virus OKIAV246 | QMP82218.1 | Bundaberg bee virus 1 | AWK77854.1 |
| Aphid bunyavirus 1 | UZM08148.1 | Planococcus ficus-associated dicistrovirus 1 | UFZ21068.1 |
| Hangzhou frankliniella intonsa phasmavirus 1 | UHK03329.1 | Leibnitzia anandria dicistrovirus | QKF95572.1 |
| Guagua virus | YP_010840748.1 | XiangYun picorna-like virus 3 | UUG74228.1 |
| Sanya sesamia inferens phasmavirus 1 | UHR49914.1 | Phenacoccus solenopsis virus | QIU80542.1 |
| Sanxia Water Strider Virus 2 | YP_010085075.1 | Anagyrus vladimiri dicistrovirus | UZC83387.1 |
| Wuhan Insect virus 2 | YP_009270651.1 | Diversinervus elegans virus | QQD86568.1 |
| ***(C) Mesoniviridae*** |  | Soybean thrips picorna-like virus 1 | QQP18805.1 |
| Aphis citricidus meson-like virus | YP_010800405.1 | Renmark bee virus 1 | AWK77866.1 |
| Yichang virus | QPK41102.1 | Anagyrus vladimiri iflavirus | UZC83388.1 |
| Frankliniella occidentalis associated mesonivirus 1 | YP_010800379.1 | Planococcus ficus-associated iflavirus 1 | UFZ21065.1 |
| Ofaie virus | YP_009666297.1 | Tetrastichus brontispae RNA virus 3 | UIX27085.1 |
| Alphamesonivirus daknongense | YP_007697636.1 | Wuhan coneheads virus 1 | YP_009342053.1 |
| Bontang virus | AGL73177.1 | **(F) *Virgaviridae*** |  |
| Kadiweu virus | YP_009666293.1 | Barley aphid RNA virus 3 | BBV14751.1 |
| Wuhan nido-like virus 1 | YP_009345058.1 | Drosophila River Almond nege-like virus | WPV74320.1 |
| Wuhan insect virus 19 | YP_009342322.1 | Aphis glycines virus 3 | UTQ79664.1 |
| Bemisia tabaci nido-like virus 1 | QWC36473.1 | Hubei Wuhan insect virus 9 | YP_009337901.1 |
| Fuefuki virus | AWA82245.1 | Barley aphid RNA virus 4 | BBV14754.1 |
| Nidovirales sp. | URA30378.1 | Insect virga-like virus 1 | QNM37814.1 |
| **(D) *Flaviviridae*** |  | Bemisia tabaci virga-like virus 2 | QWC36453.1 |
| Sitobion miscanthi flavi-like virus 1 | BBV14761.1 | Virgaviridae-like virus 5 | WPR18886.1 |
| Macrosiphum euphorbiae virus 1 | YP_009175071.1 | Cotesiavirus virgavi | DBA44328.1 |
| Orthopteran flavi-related virus | QTJ63573.1 |  |  |
| **(G) *Tombusviridae*** |  | Scaphoideus titanus toti-like virus 1 | QIJ56906.1 |
| Tomato associated tombus-like virus 1 | UTQ50978.1 | Sanya nephotettix cincticeps totivirus 1 | UHK03214.1 |
| Golovinomyces cichoracearum associated tombus-like virus 1 | QLL27731.1 | Circulifer tenellus virus 1 | YP_003800003.1 |
| Korle-bu Aedes virus | BBN50999.1 | Soybean thrips-associated dsRNA virus-1 | QQP18723.1 |
| Soybean thrips tombus-like virus 5 | QQP18795.2 | Sogatella furcifera toti-like virus 2 | WGU15425.1 |
| Beihai tombus-like virus 11 | YP_009337762.1 | Hangzhou totivirus 12 | UHK03335.1 |
| Sanya tombus-like virus 1 | UHK03141.1 | **(I) Reoviridae** |  |
| Hangzhou tombus-like virus 1 | UHK03273.1 | Planococcus ficus-associated reovirus 1 | UFZ21074.1 |
| Hymenopteran tombus-related virus | QTJ63634.1 | Zoersel tick virus | QYV43119.1 |
| Hydrocotyle vulgaris tombusvirus | QKF95567.1 | Shelly beach virus | AYP67577.1 |
| **(H) *Totiviridae*** |  | Operophtera brumata reovirus | YP_392501.1 |
| Bemisia tabaci toti-like virus 1 | QVT76732.1 | Eccles virus | AWA82237.1 |
| Hangzhou nephotettix cincticeps totivirus 1 | UHK03282.1 | High Island virus | AVO64750.1 |
| Spissistilus festinus virus 1 | YP_003800001.1 | Soybean thrips-associated dsRNA virus 3 | QQO81407.1 |
| Ips spici-like virus 1 | WPV71137.1 | Reoviridae sp. | WWV87442.1 |

